# The Proline-rich Domain Promotes Tau Liquid Liquid Phase Separation in Cells

**DOI:** 10.1101/2020.05.04.076968

**Authors:** Xuemei Zhang, Michael Vigers, James McCarty, Jennifer N. Rauch, Glenn H. Fredrickson, Maxwell Z. Wilson, Joan-Emma Shea, Songi Han, Kenneth S. Kosik

## Abstract

Tau protein *in vitro* can undergo liquid liquid phase separation (LLPS); however, observations of this phase transition in living cells are limited. To investigate protein state transitions in living cells we found that Cry2 can optogentically increase the association of full lengh tau with microtubules. To probe this mechanism, we identified tau domains that drive tau clustering on microtubules in living cells. The polyproline rich domain (PRD) drives LLPS and does so under the control of phosphorylation. These readily observable cytoplasmic condensates underwent fusion and fluorescence recovery after photobleaching consistent with the ability of the PRD to undergo LLPS *in vitro*. In absence of the MTBD, the tau PRD co-condensed with EB1, a regulator of plus-end microtubule dynamic instability. The specific domain properties of the MTBD and PRD serve distinct but mutually complementary roles that utilize LLPS in a cellular context to implement emergent functionalities that scale their relationship from binding alpha-beta tubulin heterodimers to the larger proportions of microtubules.

## Introduction

Many proteins have domain structures in which a contiguous amino acid sequence can operationally serve a discrete function, but in the context of the entire protein become regulated within a larger sequence space. The domain structure of tau is well recognized but the functions of these domains, and particularly the internal control they exert over tau function, are poorly understood. Tau protein undergoes a liquid to solid phase transition in numerous neurodegenerative conditions, most prominently Alzheimer’s disease, frontotemporal dementia and chronic traumatic encephalopathy as well as more rare conditions such as Pick’s disease, progressive supranuclear palsy, corticobasal degeneration, Neiman-Pick type 3, and sub-acute sclerosing encephalitis among others (Wang and Mandelkow 2016). In the past few years, liquid liquid phase separated states called condensates in cells have become associated with several proteins that accumulate in neurodegeneration including FUS (Patel et al. 2015, Maharana et al. 2018), TDP-43 (Gasset-Rosa et al. 2019, Mann et al. 2019), HNRNPA1 (Lin et al. 2015, Gui et al. 2019), and DDX (Nott et al. 2015), as well as tau (Zhang et al. 2017). An important feature of condensates is the very high concentration of protein within a small volume compared to the solute phase, while the protein constituents of these compartments are hydrated and freely exchange. The thermodynamics of liquid liquid phase separation (LLPS) of biopolymers from solution to a condensed state has been studied extensively using polymer physics models and tools (Brangwynne, Tompa and Pappu 2015, Lin et al. 2019), but the correlates of these events in living cells are mostly unknown. LLPS *in vitro* can be well-described as simple or complex coacervates; however, condensates in living cells involve multiple diverse components whose binding affinities and concentration are dynamically regulated by an “*in vivo* phase diagram” with a staggering complexity of phases and intrinsic variables that control the transitions among them. While *in vitro* studies can prove the sufficiency of molecules to undergo LLPS they strip the system of its inherent complexity. A complementary technique for grasping the function of phase transitions, and how they are regulated *in vivo* is to make acute changes in the valency of a core scaffolding protein by fusing it to a photo-sensitive oligomerization domain, thereby gaining the ability to abruptly alter the location in phase space within the cytoplasm.

One way to achieve this light-mediated change in protein valency is to use the optogenetic protein Cry2, the photolyase homology region (PHR) of a cryptochrome from *Arabidopsis thaliana*, that undergoes light-sensitive self-association upon blue light exposure (Taslimi et al. 2014). Cry2WT-mCherry (Cry2 wild type) by itself will not undergo LLPS in heterologous cells, but when fused to the IDR (intrinsically disordered region) of FUS, HNRNPA1 or DDX4 these proteins can drive droplet formation (Shin et al. 2017). LLPS of these proteins likely involves additional cellular factors that lower the free energy for LLPS that the blue light activated Cry2 mimics. Tau is an intrinsically disordered protein (IDP), binds RNA with multivalent interactions and, like the IDR domains of many RNA binding proteins (Elbaum-Garfinkle et al. 2015, Lin et al. 2015, Patel et al. 2015, Molliex et al. 2015), tau can undergo phase separation *in vitro* (Zhang et al. 2017). Here we demonstrate the potential of tau for LLPS in living cells, and identify the tau polyproline rich domain (PRD) as the regulator of condensate formation.

## Results

### Tau condensation facilitates MT binding

We cloned a full-length tau construct fused to Cry2 wild type mCherry (CWT 1-441), and in the absence of blue light activation we observed the expected microtubule (MT) pattern and some diffuse signal in the cytoplasm. With Cry2 activation by blue light, the signal from the diffuse pool was markedly reduced and the MT signal increased (Figure 1A-C). The increased tau binding to MT was quantified by computing the pixel coefficient of variation across the cell, with the lower value indicating a more uniform distribution and a higher value indicating discrete localization (Figure 1 – figure supplement A). This result suggested that tau condensate enhanced its binding to microtubules. To support this observation, we expressed Tau 1-441-EGFP (Figure 1-figure supplement B) in HeLa cells and treated cells with 1,6-hexanediol (1,6-HD) which interferes with weak hydrophobic protein-protein or protein-RNA interactions that can promote the formation of dynamic, liquid-like assemblies (Kroschwald, Maharana and Simon 2017). Three minutes after 1,6-HD treatment, the EGFP signal appeared diffuse in the cytoplasm (Figure 1-figure supplement C). The control, 2,5-HD had no effect even after 20 minutes (Figure – figure supplement D). MTs were unaffected by these treatments as indicated by 1,6-HD treatment of cells labeled with mCherry-α-tubulin (Figure 1-figure supplement E-F). Similar results were observed with Tau 256-441-EGFP (Figure 1-figure supplement E-F-G).

**Figure 1:**
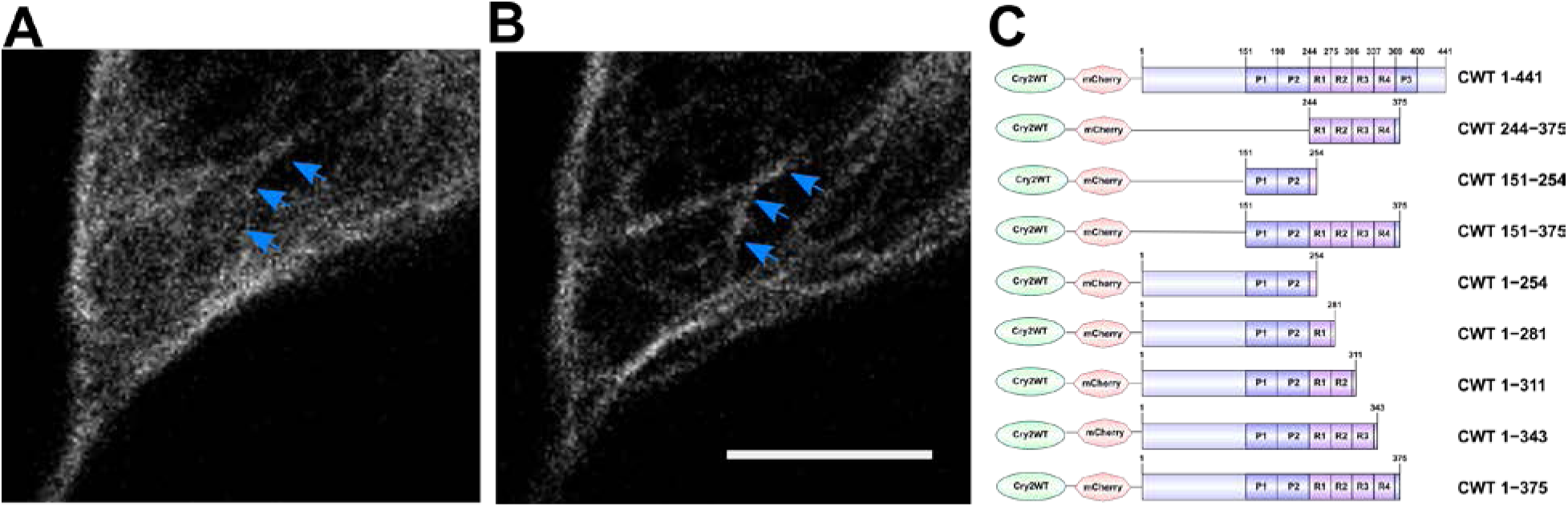
Tau condensation mediates binding to microtubules (MT). A-B) Representative Z-stack images of Cry2WT-mCherry-tau 1-441 (CWT 1-441) overexpressed in neuroblastoma SH SY5Y cell before (A) and 50 sec after activation (B). Signal from the diffuse pool was reduced and the MT signal increased. Scale Bar, 5 µm. C) Schematic diagram of Cry2WT-mCherry-tau constructs studied, the Cry2WT fused to mCherry and various tau fragments.

### The Tau PRD mediates tau condensation in cells

With the finding that tau phase separation as a condensate can enhance its binding to MTs, we sought to determine which domains of tau drive condensation-assisted MT binding and began by analyzing them individually. The tau domains consist of: the amino (N)-terminal domain, the proline-rich domain (PRD) subdivided as P1 and P2 with ~25% prolines, the microtubule-binding domain (MTBD) with four repeat regions, and the carboxy (C)-terminal domain from amino acid 369 to 441, which includes a short P3 region flanking the MTBD (Gustke et al. 1994, Mukrasch et al. 2005) (Figure 1C). Athough the MTBD can undergo liquid-liquid phase separation (LLPS) *in vitro* (Zhang et al. 2017, Ambadipudi et al. 2017) when this domain was fused to Cry2 wild type mCherry (designated CWT 244-375) and transfected into SH SY5Y cells, blue light did not induce the cytoplasmic condensates characteristic of LLPS. In contrast, the PRD domain fused to Cry2 Wild Type mCherry (designated CWT 151-254) expressed in SH SY5Y underwent condensation within 5 sec of blue light activation (Figure 2A-B). Cry2 wild type mCherry (designated Cry2WT) alone expressed in SH SY5Y cells at a comparable concentration did not cluster following blue light activation (Figure 2B) (Taslimi et al. 2014, Lee et al. 2014). The extent of light activated condensation of PRD was quantified by computing the pixel coefficient of variation, which evaluates the overall distribution across the cell (Figure 2B).

**Figure 2:**
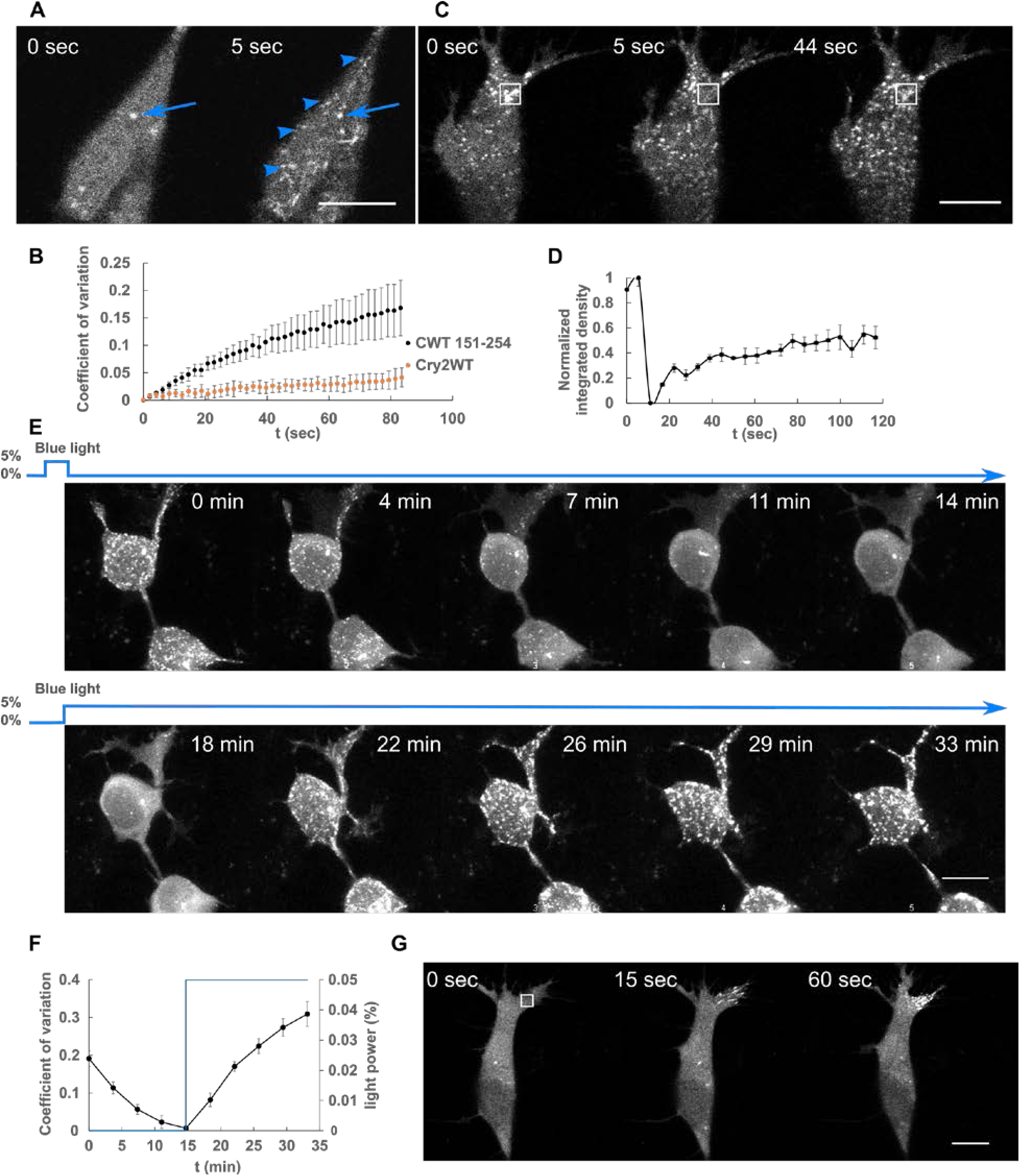
Liquid-liquid phase separation of Cry2WT-mCherry-tau 151-254 (CWT 151-254). A) CWT 151-254 overexpressed in SH SY5Y cells activated by shallow blue light (488 nm, 5% power) rapidly induced condensates (blue arrow heads). Representative fluorescent images at 0 and 5 sec after activation are shown. Occasional pre-activation foci are sometimes present and do not participate in light-activated phase separation (blue arrow). B) Temporal evolution of phase separation of CWT 151-254 (black points) and Cry2WT (orange points) were monitored on a single confocal plane, quantified by coefficient of variation across the cell and plotted over time. n = 7 for CWT 151-254 and n = 5 for Cry2WT, error bar in S.E. C) and D) Fluorescence recovery after photobleaching (FRAP) of CWT 151-254. C) Representative time-lapse images of FRAP indicated by the 3 × 3 µm bleach site is marked by the white square. See Movie S1. D) Integrated intensity density for the bleached areas were monitored over time during FRAP. Fifty-five percent recovery was achieved after bleaching. n = 3. E) and F) CWT 151-254 phase separation is reversible. E) Time-lapse images of Maximum Intensity Z-projection of CWT 151-254 shows phase separation is reversible after the withdraw of blue light (0-14 min) and reassembled when subject to blue light reactivation (14-33 min). F) Quantification of CWT 151-254 from disassembly to reassembly as light power changes. CWT 151-254 distribution quantified by coefficient of variation (black) and light activation indicated by the blue light power (blue) were ploted over time. n = 8, error bar in S.E. G) Time-lapse images of local shallow light activation of CWT 151-254 demonstrate localized progression toward phase separation. A rectangle area of 3 × 3 µm (white square) was stimulated with blue light. See also Movie S2 for local activation and the field activation followed. Scale Bar, 10 µm.

To probe the bulk exchange dynamics of light-induced CWT 151-254 condensates, we performed fluorescence recovery after photobleaching (FRAP) by bleaching the mCherry signal (Figure 2C-D, Movie S1). CWT 151-254 can exhibit up to 55 ± 15% (n = 3) fluorescence recovery. Droplets formed by blue light activation of CWT 151-254 disassembled within a few minutes in the absence of blue light, then reassembled when exposed again to blue light (Figure 2E-F) indicating that droplet formation of this construct is initially reversible. Local activation of CWT 151-254 by directing the light to a 3 × 3 µM square demonstrated that phase separation can be spatially localized (Figure 2G, Movie S2 for local field activation). As expected for light activated phase separation, tau condensation likely involves weak multivalent interactions that can be strengthened and aided in their detection by Cry2 (Bracha, Walls and Brangwynne 2019). Taken together, these data show that the PRD can undergo light-induced condensation independently of the MTBD.

We sought to determine the reversibility of tau PRD condensates over longer time periods by cycling through multiple illumination periods. A sequence of blue light exposure followed by a recovery phase with three activation cycles was performed (Shin et al. 2017) (Figure 3A). After the first cycle, the condensates were mostly dissolved, but residual condensates increased progressively after each cycle. The result was quantified by counting the condensates and normalizing the result (see methods). The percentage of condensates that remained was 0%, 42%, 73% after the first, second and third cycle respectively (Figure 3B). The kinetics of cluster assembly and disassembly were also analyzed by pixel coefficient of variation (Figure 3C). As expected, the values increased during assembly and decreased during disassembly. Through the recovery, the rate of the disassembly gradually slowed and eventually reaches a plateau. Upon each cycle the recovery plateaued progressively earlier and further from the initial state. These findings resemble an “aging” process in which the liquid state transitions to a non-exchangable gel or a solid phase (Shin et al. 2017).

**Figure 3:**
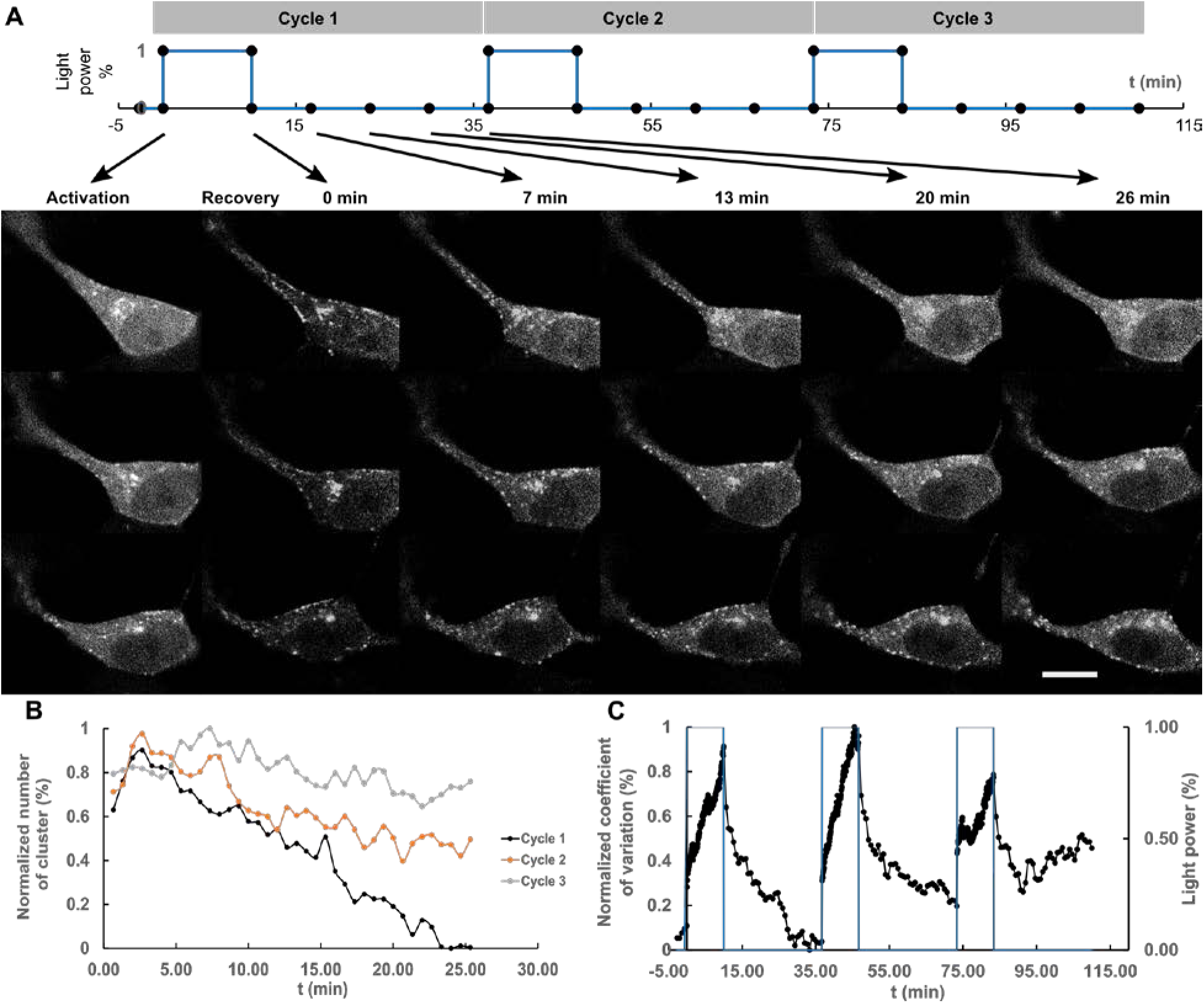
Repeated deep blue light activation reduced recovery of Cry2WT-mCherry-tau 151-254 (CWT 151-254) biocondensates. A) Top: a sequence of three cycles deep blue light activation, 488 nm, 100% power for 10 min, each followed by 26 min of recovery time applied to SH SY5Y cells, in which CWT 151-254 was overexpressed. Bottom: time lapse of single Z-stack images shows CWT 151-254 biocondensate assembly upon light activation and disassembly when light was removed following the sequence on the top. For each cycle, the image at initial activation and recovery at 0, 7, 13, 20, 26 min are shown. B) The number of the clusters post-activation was counted and plotted. C) Re-distribution of CWT 151-254 through the time course is quantified by coefficient of variation and change in light power over the time course. Scale Bar, 10 µm.

The PRD is the most heavily phosphorylated region in tau and its phosphorylation is controlled by numerous kinases in diverse signaling pathways (Martin et al. 2013). Phosphorylation in this region will modulate the charge distribution and affect the weak electrostatic interactions that mediate condensation. Placing phosphomimetic mutations within the PRD completely abolished blue light induced phase separation of CWT (Table 1). Mono-, di- or triple phosphomimetic mutations at T231E or S235E, T231E/S235E or T212E/S214E/T217E, or S202D/T205D/S208D all abolished light induced phase separation. In contrast, LLPS of CWT 151-254 with alanine mutations at the same phosphorylation sites was not abolished. Upon light activation, CWT 151-254 S202A/T205A/S208A and CWT 151-254 S202A/T205A/S208A/T212A/S214A still phase separated.

**Table 1:** Phosphomimetics in PRD as the regulation of condensation*.

| Construct | LLPS |
| --- | --- |
| CWT 151-254 | Yes |
| CWT 151-254 T231E | No |
| CWT 151-254 S235E | No |
| CWT 151-254 T231E/S235E | No |
| CWT 151-254 T212E/S214E/T217E | No |
| CWT 151-254 S202D/T205D/S208D | No |
| CWT 151-254 S202A/T205A/S208A | Yes |
| CWT 151-254 S202A/T205A/S208A/T212A/S214A | Yes |

\*All experiments were performed at the same expression levels and at constant light levels.

### PDR condensates promote Tau association with MTs

Having characterized the isolated domain behavior of the PRD, we next sought to determine its relationship to the microtubule and its behavior in the context of the neighboring MTBD. Light induced PRD condensates (CWT 151-254) were frequently observed to align along MTs (Figure 2A and Figure 4A). The MTBD on its own (CWT 244-375) and transfected into SH SY5Y cells did not bind MTs nearly as well as full length tau (Figure 4B-E). In contrast, CWT 151-375 (Figure 4F-G) which contains both the PRD and the MTBD, bound to microtubles similarly to full-length tau (CWT1-441) before activation and increased its binding further after light activation, also similarly to full-length tau (Figure 4B-C).

**Figure 4:**
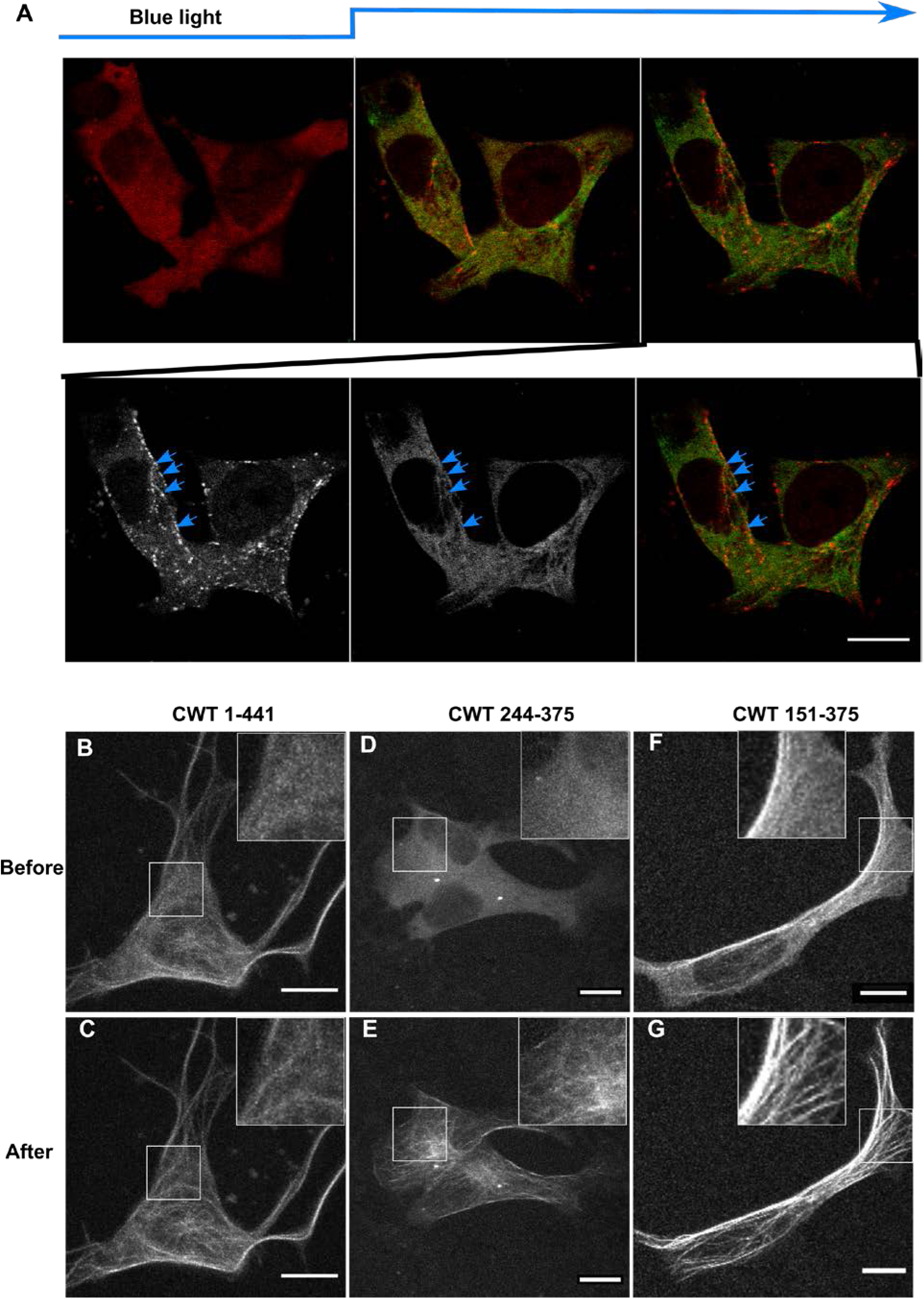
Cry2WT-mCherry-tau fragments associate with MT. A) Association of PRD condensates with microtubules. EGFP-α-tubulin (green) and CWT 151-254 (red) were coexpressed in neuroblastoma SH SY5Y cells. Time-lapse images show CWT 151-254 forms LLPS upon blue light activation starting from the 2nd time frame. Montage of two fluorescent channels in 3rd time frame demonstrates CWT 151-254 foci are attached to EGFP-α-tubulin on MT, like beads on a thread (blue arrow). B-G) Cry2WT-mCherry fused to MTBD without and with PRD were overexpressed in SH SY5Y cells and signal distribution was studied before and after blue light. Representative maximum intensity Z-projection images before and after blue light of each construct are shown, CWT 1-441 before (B) and after (C), CWT 244-375 before (D) and after (E), CWT 151-375 before (F) and after (G). Insets within the white rectangle show more details of the diffuse pool and the MT-binding population in each construct. Scale Bar, 10 µm.

Inclusion of the N-terminal domain with the PRD (CWT 1-254) had an inhibitory effect over tau condensation. Unlike the PRD domain alone, this construct showed only minimal association with MTs both before and after light activation (Figure 4-figure supplement A-B). This result suggested that the N-terminal 1-150 region negatively regulates PRD phase separation likely owing to its many negatively charged residues and is consistent with the negative regulation of N-terminal region on PRD binding to soluble tubulin and slowing polymerization (McKibben and Rhoades 2019) and previous work showing that removal of the n-terminal region increased the affinity of tau for microtubules (Gustke et al. 1994).

As additional MT binding domains were sequentially added to CWT 1-254 construct, MT binding increased and the diffuse labeling decreased (Figure 1C, Figure 4-figure supplements C-J and K-L and Figure 4-table supplement and methods). A construct from the amino terminus that included all of the MT binding repeats (CWT 1-375) further increased association with MTs after blue light, but was still not maximized until we used a full-length construct that included the P3 (376-400) domain (Figure 4-figure supplement I-J). The P3 domain flanks the MTBD and also undergoes proline-directed phosphorylation (Mukrasch et al. 2005). These data demonstrate that condensation of the PRD alone can drive association with MTs and that association is inhibited by the N-terminal domain but enhanced by the MTBD as well as the C-terminal domain.

### The tau PRD phase separates with EB1

Plus end–tracking proteins (+TIPs) form a complex interaction network at microtubule plus ends, where they control microtubule dynamics and microtubule anchorage to subcellular targets such as vesicles and kinetochores (Galjart 2010). End-binding protein 1 (EB1) is a major integrator of the +TIP complex that appears as “comets” at MT ends (Akhmanova and Steinmetz 2010). Tau forms a complex with EB1 through an interaction between the MT-binding repeats and the P2 region of the PRD interacting with the acidic C-terminal tail of EB1 (Ramirez-Rios et al. 2016). The observation that CWT 151-254 moved along MTs as a comet shaped structure during the initial 30 sec (Figure 2G), before it formed a condensate, prompted us to study the interaction of CWT 151-254 with EB1. We co-expressed CWT 151-254 and EB1-EGFP in SH SY5Y cells activated by blue light. Kymograph tracking the fluorescent “comets” (Mimori-Kiyosue, Shiina and Tsukita 2000) of EB1-EGFP showed spatiotemporal co-localization with CWT 151-254 after blue light activation in which light activation first induced comet formation that rapidly extended along MTs (Figure 5A-B, Movie S3).

**Figure 5:**
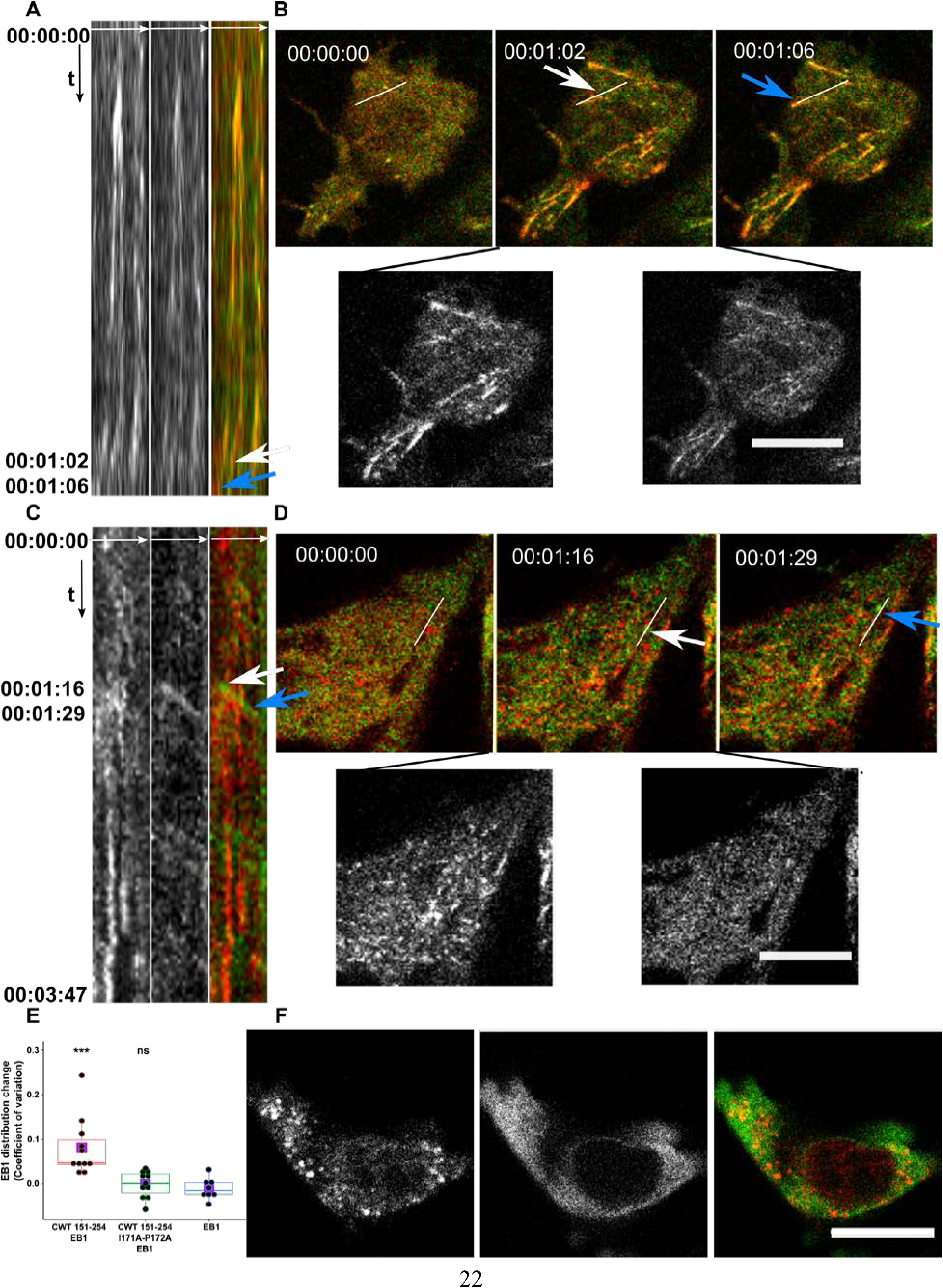
Cry2WT-mCherry-tau 151-254 (CWT 151-254) tracks with EB1 after blue light activation. A-B) CWT 151-254 overexpressed with EB1-EGFP in SH SY5Y cells and activated by shallow blue light (488 nm, 5% power) induced colocalization. A) Kymograph of CWT 151-254 (red) and EB1-EGFP (green) is generated from time lapse imaging for 66 sec at 2 sec imaging intervals, with the x axis the selected line crossing the image and the y axis the time of recording. Fluorescence imaging of two channels shows the formed CWT 151-254 (red) condensate colocalized with moving EB1-EGFP (green). B) Representative time lapse images at the time of 00:00:00, 00:01:02 and 00:01:06 are shown in the top panel, where the line used to generate the kymograph is shown in white. The kymograph in (A) moves from left to right. Selected moving foci on the line are indicated with arrows white or blue on both the kymograph and the corresponding time lapse images. Two fluorescent channels of the middle image at 00:01:02, CWT 151-254 (red left), EB1-EGFP (green, right), are presented at the bottom panel. See also Movie S3. C-D) CWT 151-254 with I171A and P172A double mutant and EB1-EGFP overexpressed in SH SY5Y cells activated by shallow blue light (488 nm, 5% power). The induced double mutant CWT 151-254 (red) condensate does not colocalize with EB1-EGFP. C) Kymograph of CWT 151-254 with I171A and P172A double mutant (red) and EB1-EGFP (green) generated from a time lapse imaging for 3 min and 47 sec at 2 sec imaging intervals, with the x axis being the selected line crossing the image and the y axis the time of recording. Fluorescence imaging of two channels shows the formed CWT 151-254 with I171A and P172A double mutant (red) condensate does not colocalize with moving EB1-EGFP (green). D) Representative time lapse images at the time of 00:00:00, 00:01:16 and 00:01:29 are shown in the top panel, where the line used to generate the kymograph is shown in white. Selected moving foci on the line are indicated with arrows white or blue on both the kymograph and the corresponding time lapse images. Two fluorescent channels the middle image at 00:01:16, CWT 151-254 I171A and P172A (red, left), EB1-EGFP (green, right), are presented at the bottom panel. E) Quantification of EB1-EGFP distribution after light activation. EB1 was expressed in SH SY5Y cells with CWT 151-254, or with CWT 151-254 I171A/P172A double mutant, or with no co-overexpression. Distribution of EB1 is evaluated by the change in the coefficient of variation after 60 sec of light activation shown along the y-axis. n = 11, 11, 8 for the groups from left to right along the x-axis. Mean for each group is indicated by a purple square. Statistical significance by pairwise t-test of each independent group against EB1 without CWT 151-254 is indicated at the top of the plot. F) CWT 151-254 (red) and EB1-EGFP (green) overexpressed in SH SY5Y cell, treated with 20 µM of nocodazole for two hours, activated by blue light induced spherical condensates. EB1-EGFP appeared diffuse in the cytoplasm and not colocalized with CWT 151-254. Representative montage image is shown. See also Movie S4. Montage of two fluorescent channels is presented from the left, the mCherry channel, the GFP channel separate and the merged. Scale Bar, 10 µm. ns, not significant; *, p < 0.05; ** p < 0.01; ***, P < 0.001.

We could not employ FRAP for these condensates because they moved too rapidly, but EB1 comets dissolved following 1,6 hexanedione treatment. Colocalized CWT 151-254/EB1 was also observed along the MT lattice (Figure 5-figure supplement) similar to EB1 when overexpressed alone (Skube, Chaverri and Goodson 2010), suggesting activated CWT 151-254 helped to concentrate EB1 on MT. The tau PRD contains one S/TxIP (169TRIP172), the consensus motif for EB1 binding (Honnappa et al. 2009). A double mutation in this consensus binding motif of tau (I171A and P172A) abolished colocalization with EB1-EGFP (Figure 5C-D). Quantification of EB1-EGFP/CWT 151-254 colocalization following light activation confirmed the presence of this complex (Figure 5E).

The EB1-EGFP/CWT 151-254 interaction was disrupted by the same phosphorylation sites that disrupted CWT 151-254 alone. Multiple mono-, di- or triple phosphomimetic mutations, T231E or S235E, or T231E/S235E or T212E/S214E/T217E, or S202D/T205D/S208D all completely abolished comets (Table 1), whereas CWT 151-254 with alanine mutations at the same phosphorylation sites did not abolish comets.

The association of the CWT 151-254/EB1 complex with MTs prompted us to examine the colocalization of the complex following treatment with nocodazole (Figure 5F, Movie S4). Under these conditions, the EB1 signal became diffuse and CWT 151-254 formed spherical condensates no longer colocalized with EB1. Thus, CWT 151-254/EB1 complexes were dependent upon the presence of polymerized MTs.

### *In vitro* tau PRD phase separation

The distinct properties of the PRD and the MTBD when fused to Cry2 prompted further investigation of these constructs *in vitro.* This comparison was especially important because we previously showed that an MTBD construct tau 255-441 can readily undergo phase separation when utilizing RNA or heparin as a cofactor (Zhang et al. 2017, Lin et al. 2020) This construct included the sequence observed in the insoluble tau fibril by cryo-electron microscopy (Fitzpatrick et al. 2017). Tau 151-254 protein with a 6xHIS tag at the N-terminus was purified. This protein was mixed with the cofactor in a 1:1 charge ratio. The occurrence of phase separation was evaluated by measuring turbidity via absorbance at 500 nm, and fibril formation measured by fluorescence of Thioflavin T (ThT) that binds to beta-sheets of stacked tau proteins. Directly after addition of polyU to tau 151-254, droplets formed (Figure 6A) that were capable of fusing and deforming as they wet the glass surface. Tau 255-441 mixed with the heparin cofactor not only formed droplets which fused and deformed as they wet the glass surface, but also induced significant ThT fluorescence (Lin et al. 2020). In contrast to tau 255-441, tau 151-254 did not show any ThT fluorescence (Figure 6B). TEM imaging confirmed that no fibrils were formed by the tau 151-254. Although both tau 151-254 and tau 255-441 readily formed droplets *in vitro* with the addition of the negatively charged cofactors, polyU or heparin (Figure 6A), condensation was insufficient for the PRD to make fibrils. This distinction may be related to energetics involved in the difference between tau 255-441 versus tau 151-254 to form fibrils.

**Figure 6:**
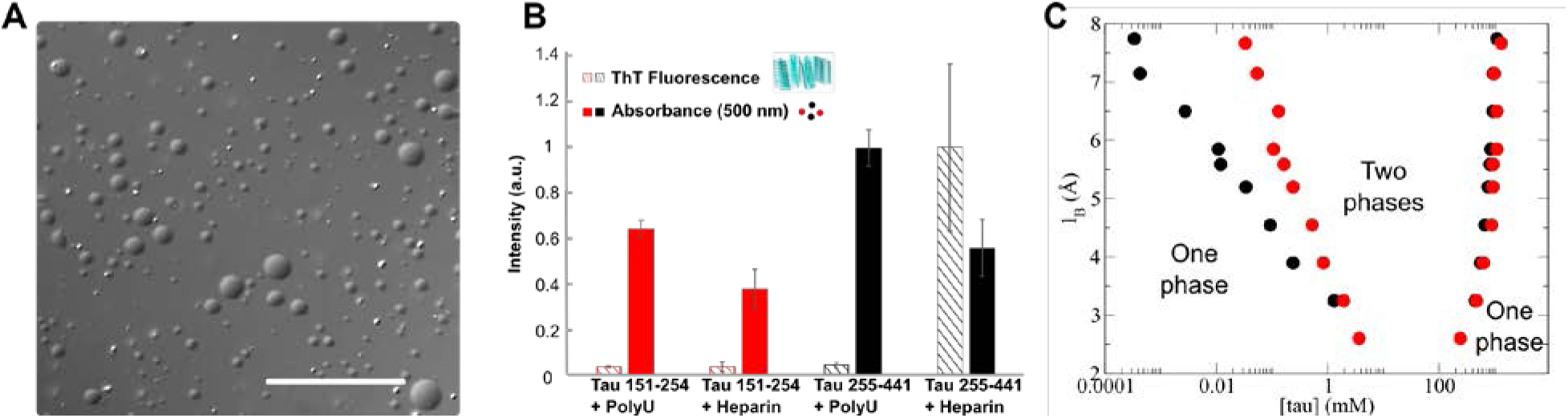
Similar to tau 255-441, tau 151-254 can form droplets *in vitro*. A) Representative bright-field images of tau 151-254/poly(U) droplets, Scale bar is 50 µm. B) Direct comparison of tau 151-254 (red) and tau 255-441(black) shows phase separation occurs for both protein segments with addition of polyU or heparin. In contrast to the tau 255-441 heparin complex, tau 151-254 does not show the capacity to form fibrils with either polyU or heparin. Turbidity was evaluated at 500 nm (filled bar), fibrilization was evaluated by Thioflavin T (ThT) fluorescence (striped bar). Protein concentration was 50 µM, polyU concentration (where applicable) was 125 µg/mL, and heparin concentration was 12.5 µM. All data were normalized by the largest measured value or absorbance or fluorescence. n = 3, error bar in standard deviation. C) Predicted tau 151-254/RNA binodal phase coexistence points modeled using field theoretic simulations (FTS). Calculations performed for tau 151-254 (red dots) are compared with our previously reported (Lin et al. 2019) phase diagram for tau 255-441 (black dots), showing that both sequences have a stable two-phase region under similar electrostatic environments and solvent conditions. The conditions for stable binodal phase coexistence depend on the strength of the electrostatic interaction, characterized in our model by the Bjerrum length (l_B_) as the input parameter. The region defined by the coexistence points is the region at which the system forms a thermodynamically stable droplet phase in coexistence with a solution phase. Areas corresponding to one-phase or two-phase are shown in the diagram.

To test if the charge patterning of the proline-rich domain (tau 151-254) leads to a similar charge-driven phase separation by the complex coacervation mechanism (Zhang et al. 2017) as compared to the MTBD (tau 255-441), we computed a phase diagram for the LLPS of a polymer physics model of tau 151-254 using a numerical method called field theoretic simulation (FTS)(Lin et al. 2019). The details of the FTS calculations can be found in the supplemental section. In our computational model, the protein is described as a bead-spring polymer with a charge sequence determined by the primary amino acid sequence of tau at pH 7.4 in presence of RNA (Figure 6C). The phase diagram depends on the strength of the electrostatic interaction, characterized in our model by the Bjerrum length (l_B_) as the input parameter which controls the strength of the Coulombic interaction according to the pair potential energy 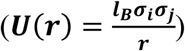. All simulations were performed in implicit solvent with statistical segments interacting through a weak excluded volume potential. Binodal points are plotted as a function of tau concentration vs. the Bjerrum length that characterizes the extent of electrostatic screening.

At low l_B_, the electrostatic strength in the model is not sufficient for phase separation, but at l_B_ > 2.5 Å our model of both tau 151-254 and tau 255-441 have a region where phase separation occurs (Figure 6C). The left points show the lower concentration limit for a stable droplet in solution, and the right points show the upper concentration window. We found that tau 255-441 and tau 151-254 have a similar propensity to form droplets, as manifest by a similar phase diagram (Figure 6C). As a negative control, both the N-terminal region of tau (tau 1-150) and the C-terminal region of tau (376-441) have been modeled and found not to show any stable two-phase window at these values of l_B_. Thus, the biophysical properties of the tau PRD and MTBD are similar in term of their ability to phase separate; however, their differences are revealed in the biological context of a living cell.

## Discussion

The discovery that tau is capable of LLPS *in vitro* (Zhang et al. 2017, Wegmann et al. 2018, Hernandez-Vega et al. 2017, Ambadipudi et al. 2017, Tan et al. 2019, Siahaan et al. 2019) raised the question of whether similar phase states of tau are present in living cells. Although both the full-length tau and MTBD are capable of phase separation *in vitro* (Lin et al. 2019, Fichou et al. 2018), when expressed in cells, these constructs did not form LLPS droplets under any of a variety of conditions. We amplified the weak multivalent interactions of tau in cells by fusing it to a light-activatable self-associating Cry2(WT) motif while ascertaining that Cry2(WT) alone did not phase separate under the same conditions. The fusion of “sticky” intrinsically disordered domains (IDR) to the photolyase homology region (PHR) of Arabidopsis thaliana Cry2 (Bugaj et al. 2013), has been used to demonstrate condensation of aggregation domains in RNA binding proteins (Shin et al. 2017). For example, the IDR of FUS fused to Cry2WT as well as HNRNPA1 or the N-terminal IDR of DDX4 (Shin et al. 2017), an IDR-Cry2 fusion protein can confer tunable light-dependence to its multivalent interactions (Shin et al. 2017). In our hands, this approach identified the unique function of the tau PRD (151-254) in driving condensation under a variety of experimental conditions. Focused blue light at specific subcellular locations can locally induce CWT 151-254 phase separation (Figure 2G), which undergoes fluorescence recovery after photobleaching (FRAP) (Figure 2C-D). The multivalent interactions of these condensates were controlled by the tau phosphorylation state (Table 1), and therefore tau condensates could be tuned by the complex phosphorylation patterns of tau. The majority of the more than 40 known tau phosphorylation sites are located in the PRD (Gong et al. 2005) and are mediated by kinases that participate in multiple signaling pathways.

Our *in vitro* and *in silica* data support that the PRD can form droplets with the addition of negatively charged cofactors similar to the microtubule binding domain (MTBD). Interestingly, the experimental and theoretical biophysics show that the LLPS propensity of MTBD and PRD are indistinguishable, and therefore, their difference lies in their biological properties. Among these properties is complex multiple phosphorylation of the PRD which results in vastly more charge variations to work in concert with the MTBD.

Considerable interest has mounted in the detailed relationship of tau to MTs. Cryo-electron microscopic images of the tau microtubule-binding domains demonstrate an interaction with the MT surface on which single repeats bind across an alpha-beta tubulin dimer (Kellogg et al. 2018). In a surprising finding, McKibben and Rhoades (McKibben and Rhoades 2019) showed that the isolated PRD bound tubulin tightly compared to the microtubule binding domain, exhibited a higher affinity for taxol-stabilized microtubules, and was the only isolated domain capable of significantly polymerizing tubulin. These authors found that under saturable stoichiometric binding conditions one tau bound to two tubulin dimers. Therefore, based solely on the observations in both studies, the number of tau molecules that bind to MTs is in the range of 400 to 800 copies within a one um segment of MT given a 13 protofilament structure with ~1600 alpha/beta tubulin heterodimers (Löwe et al. 2001). However, it is widely agreed that numerous factors will alter the number of bound tau molecules including tau splicing in the repeat region, the state of tau phosphorylation and acetylation, the specific locus of binding along the MT lattice, and the tubulin concentration among others. Despite the difficulty of determing precise numbers of tau molecules at the scale of polymerized MTs in cells, certainly hundreds or thousands of tau molecules are concentrated within a small volume for MT polymerization and stability. Phase separation provides high local concentrations of tau exceeding that of the solute phase in which molecular collisions would make precisely oriented tau-MT binding improbable. Rather tau proteins can collectively implement their function at the larger scale of MTs by securing the assembly of multiple alpha-beta tubulin dimers nearly instanteously.

McKibben and Rhoades (McKibben and Rhoades 2019) suggest that the PRD serves as the core tubulin binding domain, binding to two tubulin dimers as a critical step towards initiating polymerization, and multiple weak tubulin binding sites in the MTBD increase the local concentration of tubulin to accelerate microtubule growth. Thus, the differential properties of these two domains ultimately lead to the assembly and stability of a MT. Unfortunately, the PRD was not visualized in the cryo-electron microscopy of the tau/MT interaction (Kellogg et al. 2018)). Tan et al (Tan et al. 2019) used bacterially-expressed, GFP-tagged full-length human tau to bind to taxol-stabilized MTs *in vitro*. In this study, tau molecules initially bound diffusely along the entire MT lattice and progressed to equilibrium as condensates. Interestingly, tau condensates had increased average dwell time along the MT lattice (Tan et al. 2019), thereby taking advantage of condensate/solute phase equilibrium properties that will sustain a high local concentration of tau in proximity to the MT. Phase separated droplets can also regulate their interfacial tension and thereby wet the solid MT surface.

Tan et al (2019) also show that tau condensation only formed on GDP MT lattices suggesting that tau condensation was gated by the nucleotide state of the MT lattice and therefore does not condense in the region of the GTP-Cap. The inhibition of tau condensation at the GTP-Cap may function in conjunction with the co-condensation of tau CWT 151-254 and EB1, that we observed. EB proteins control the protofilament number, the length of taper at microtubule tips and mechanically stiffen the microtubule (Vitre et al. 2008, Lopez and Valentine 2014, Zhang et al. 2015) as well as track the ends of growing microtubules (Bieling et al. 2007, Bieling et al. 2008, Komarova et al. 2009) by recognising a nucleotide-dependent conformation is transiently formed during microtubule assembly of two adjoining tubulin subunits (Roth et al. 2019). Exchange between CWT 151-254/EB1 complexes and tau condensates on the MT lattice may coordinate the nucleotide state of the MT with its polymerization capacity within the region of the EB comet.

In a living cell, proteins retain many of their fundamental biophysical properties that become highly regulated and specified to achieve precision functions in the context of numerous dynamic conditions. The PRD has a prominent role in regulating tau LLPS in living cells, a function that we detected by combining *in vitro* and *in vivo* studies along with the optogenetic probe, Cry2. Tau condensates mediated by the PRD can create finely tuned molecular locales to regulate cellular architectures under the control of MTs.

### Contact for Reagent and Resource Sharing

Further information and requests for reagents may be directed to Kenneth S. Kosik.

### Experimental Model and Subject Details

#### Reagents

Nocodazole and PolyU (RNA) (MW 800~1000 KDa) were purchased from Sigma, 1,6-and 2,5-Hexanediol from Aldrich, Heparin from Galen Laboratory Supplies.

#### Cell culture

SH-SY5Y neuroblastoma cells (ATCC, CRL-2266) in DMEM/F12/GlutaMAX medium (Gibco) with 10% FBS or Hela cell in DMEM medium with 10% FBS were cultured as monolayers at 2 × 10^4^ per well on 8-well µ-Slides (ibidi) at 37°C with 5% CO_2_ in a humidified incubator. The population of neuroblast-like cells may decrease with increased passage number (Kovalevich and Langford 2013); therefore, the passage number was kept below 30 and differentiation induced by nutritional deprivation was carefully avoided.

## Method Details

### Plasmid and recombinant protein

#### Plasmids for mammalian expression

Plasmid Cry2olig-mCherry-tau 1-441 was prepared by inserting DNA fragment encoding the full length tau into the linearized Cry2olig-mCherry (Addgene 60032) backbone at the C-terminus of mCherry using Gibson assembly® Cloning kit (New England BioLab Int.). Cry2WT-mCherry-tau 1-441 is then produced by reversing the E490G mutation of Cry2olig. Plasmids Cry2-mCherry-truncated tau including Cry2WT-mCherry-tau 244-375,151-254,1-375, 1-343, 1-311,1-281, 1-254, 151-375 were prepared from Cry2WT-mCherry-tau 1-441 using Q5 Site-Directed Mutagenesis Kit (New England BioLab Int.). Plasmids Cry2WT-mCherry-tau 151-254 psuedo-phosphorylated mutants or alanine substitutes were prepared from Cry2WT-mCherry-tau 151-254 using the same mutagenesis kit. Plasmid Cry2olig-mCherry-tau 151-254 was created in the similar way from Cry2olig-mCherry-tau 1-441. Unless otherwise mentioned all the transformation were done in high efficient competent cell (New England BioLab Int.) and all the resulting plasmids were partially sequenced to confirm the coding sequence. Tau 1-441-EGFP and Tau 256-441 EGFP were gifts from Dr. Gerold Schmitt-Ulms, at University of Toronto, Ontario, Canada. The plasmids encoding EGFP-α-tubulin (addgene 12298), pmCherry-α-tubulin (Addgene 21043) and EB1-EGFP (Addgene 39299) were purchased.

#### Plasmids for bacterial expression and recombinant proteins

cDNA of tau 151-254 was inserted into a pMCSG17 plasmid using HiFi assembly (NEB #E2621S). The resulting vector encoded a protein consisting of a hexahistidine tag followed tobacco etch virus (TEV) enzymatic cleavage site attached to the tau 151-254 sequence. The plasmid was transformed into DH5a, and BL21DE3 cells.

Tau 255-441 was purified as previously described (Zhang et al. 2017). A similar procedure was used for purification of Tau 151-254, however the 65°C heatshock stage of purification was not used. Following Ni-NTA affinity purification, size exclusion chromatography was used. The elution fraction from the Ni-NTA was collected and concentrated, then loaded onto a Biorad ENrich 70 SEC column pre-equilibrated with 20 mM HEPES and 40 mM NaCl buffer. Fractions were collected and pooled based on UV absorption at 280 nm. The species identity was confirmed with SDS PAGE, and MALDI-TOF mass spectroscopy. The concentration was determined with a bicinchoninic acid (BCA) assay, or from the absorption at 280 nm.

### Live cell imaging

Cells were plated on the No. 1.5H 170 µm glass-bottom 8-well µ-Slides (ibidi) and grown typically overnight in normal growth medium to reach ~ 50% confluency. Live cell imaging was performed using 60X glycerol immersion objective on a Leica Sp8 laser scanning confocal microscope equipped with a temperature stage at 37°C and suppled with 5% CO_2_. Cells were imaged typically using two laser wavelengths (488 nm for Cry2 activation and GFP channel imaging /555 nm for mCherry) and sequentially scanned by line. For activation, unless specifically mentioned, three 488nm light power of 5%, 30% and 100% were used for shallow, intermediate and deep activation condition respectively. Cell was monitored over time in either xyt or xyzt mode. For local activation of CWT 151-254, a square of 3 × 3 µm was stimulated with blue light, and the dynamics of the region quantified.

For fluorescence recovery after photobleaching (FRAP), cells were first activated by 488 nm with an appropriate blue light intensity. Immediately after termination of the activation phase, light-induced clusters were bleached by UV light with 100% power on a square of 3 x3 μm for a period of time, depending the signal intensity of site, and their fluorescence recovery was monitored while maintaining identical activation conditions for the clustering. It is practically helpful to bleach the site to the point that it is still noticeable, so the signal recovery can be precisely followed up.

### Live cell imaging analysis

#### Phase separation quantified by single cell coefficient of variation of labeled proteins

Phase separation results in non-uniform distributions of phase separated proteins in cells. When labeled proteins are diffusely distributed their coefficient of variation (CV), i.e. the standard deviation in pixel intensity from all measured points in the cell over the mean signal across the entire cells, is an excellent measure of the non-uniformity. The images were analyzed in Fiji-ImageJ.

#### Fluorescence recovery after photobleaching

Integrated intensity density of the area for single Z-stack was monitored, corrected for photobleaching with a reference area of the same size and normalized with pre-bleaching intensity.

#### Reversibility of clustering

Z-projection of the whole cell, applying the standard deviation method embedded within imageJ, was monitored over time. The CV of the region of interest was plotted over the time course. For repeated activation and recovery experiments, images of single Z-stacks were monitored and analyzed by ImageJ by both counting the particles and by analyzing the CV of the pixel distribution in each image. To count the particles, raw images were smoothed through Gaussian Blur then the Threshold was set based on the preactivation intensity. The clusters formed after activation in each frame were counted through the imageJ function of Analyze Particles. The number of clusters in each image was then normalized according to the Max and Min number of clusters in all three cycles.

#### Analysis of tau condensation on MT binding

Cells were typically imaged by use of two laser wavelengths (488 nm for Cry2 activation /555 nm for mCherry imaging). To quantify Cry2WT-mCherry tau relocation, Z-stack imaging was performed systemically in five cycles. For the first cycle the imaging was done with only the 555 nm laser, the following four cycles with both 488nm /555nm lasers in a sequential “between lines” manner. Each cycle took 60 ± 5 sec and 10% power of the 488nm laser was used for activation. The images in the 1^st^ cycle and the 3^rd^ cycle were selected for image analysis, corresponding to Cry2 WT tau before and 60 sec after blue light activation. Images were generated from cells plated and transfected on different days.

The relocation of CWT from the diffused state to MT-binding state in a cell was monitored by CV. Custom written macros were used to automate image analysis and quantification process. Briefly, raw 3D image was first Z-projected into a 2D image applying the standard deviation method embedded in ImageJ. Each individual cell was outlined as the region of interest (ROI) with particle analyzing tools after the Z-projection 2D image was subjected to background subtraction and Gaussian filter. The CV within the ROI was then measured and reported for Cry2 WT-mCherry tau distribution before and after blue light activation in each cell.

We applied the second image analysis method to visualize tau distribution change in a cell. Using the built-in plugin “image calculator” with the operation of subtraction, the difference of the Z-projection 2D image before and after activation was calculated. With proper thresholding, a binary 2D image was generated depicting the area where the CV was increased after activation. The detected distribution shift in each cell was examined manually to validate the CV values.

### *In vitro* Turbidimetry and brightfield microscopy

Turbidity of samples at room temperature measured by optical density at a 500 nm wavelength (OD_500_), using a BioTek synergy2 plate reader. The amount of coacervates in a sample were approximated to be proportional to its maximum OD_500_. Brightfield microscopy images of condensates were taken with a Leica BX51 or IX51 compound microscope.

### *In vitro* Tau PRD and polyU condensate formation

Unless otherwise stated, condensates of tau PRD with polyU RNA or heparin were formed with 50 µM Tau PRD and 125 µg/mL polyU, and heparin concentration was 12.5 µM. This corresponds to a positive:negative charge ratio of 1:1. Ambient temperature and 20 mM HEPES, 40 mM NaCl pH7 buffer were used.

### Computational modeling of the MTBD and PRD charge pattern

The strength of the Coulombic interaction was parameterized according to the potential energy 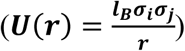, where l_B_ controls the strength of the Coulombic interaction (For reference, l_B_ = 0.71 nm for water at 300K). All simulations were performed in implicit solvent with statistical segments interacting through a weak excluded volume potential. Binodal points were plotted as a function of tau concentration versus the Bjerrum length that characterizes the extent of electrostatic screening. Simulations are performed by numerically propagating a complex Langevin (CL) equation of motion for the complex-valued fields. See supplement for detailed methods and parameterization. All simulations were performed on NVIDIA Tesla M2070 or K80 graphics processing units (GPUs) using the PolyFTS code maintained by the Fredrickson group at UCSB and licenced through the Complex Fluid Design Consortium.

### Quantification and Statistical Analysis

Statistical parameters are indicated in the legends of each figure including the definitions of error bars (standard error, S.E., or standard deviation, StDev) and the number experimental replication, denoted by n. For light activation induced change of tau distribution with each independent construct, pairwise t-test was performed.

### Data and Code Availability

Source data for figures in the paper is available upon request.

## Author Contributions

K.S.K and X.Z. conceived the project. X.Z. performed the cellular experiments. M.V. performed the *in vitro* experiments. J. M. perfomed *in silico* simulation. J.N.R. and M.W. advised with methodologies. X.Z., M.V., J.M., S.H., K.S.K wrote the manuscript. All authors edited and reviewed the manuscript. K.S.K. supervised the cellular work. S.H. supervised the *in vitro* work. J.S. and G.H.F supervised the *in silico* simulation.

## Acknowledgements

This study was funded by U54 NS 100717 (K.S.K.), Tau Consortium (K.S.K., S.H.), Dr. Miriam and Sheldon G. Adelson Medical Research Foundation (K.S.K., M.W.), Larry L. Hillblom Foundation (K.S.K.), the Edward N. & Della L. Thome Memorial Foundation (K.S.K.) and the National Institutes of Health (NIH) (grant R01AG056058 (K.S.K, S.H., J.S, J.M.). G.H.F. and J.S. acknowledge the MRSEC Program of the BSF under Award No. DMR 1720256, and J.S. the NSF under Award No. MCB-1716956. The computational part of this research used resources of the Extreme Science and Engineering Discovery Environment (XSEDE, supported by the NSF Project TG-MCA05S027) and the Center for Scientific Computing from the California NanoSystems Institute UC Santa Barbara (CNSI) available through the Materials Research Laboratory (MRL): an NSF MRSEC (DMR-1720256) and NSF CNS-1725797. We acknowledge the use of the NRI MCDB Microscopy Facility and the Resonant Scanning Confocal supported by the NSF MRI grant DBI-1625770, and Dr. Carol Vandenberg, at University of California Santa Barbara, Santa Barbara, USA, for valuable discussions.

---

**Figure 1-supplement:**
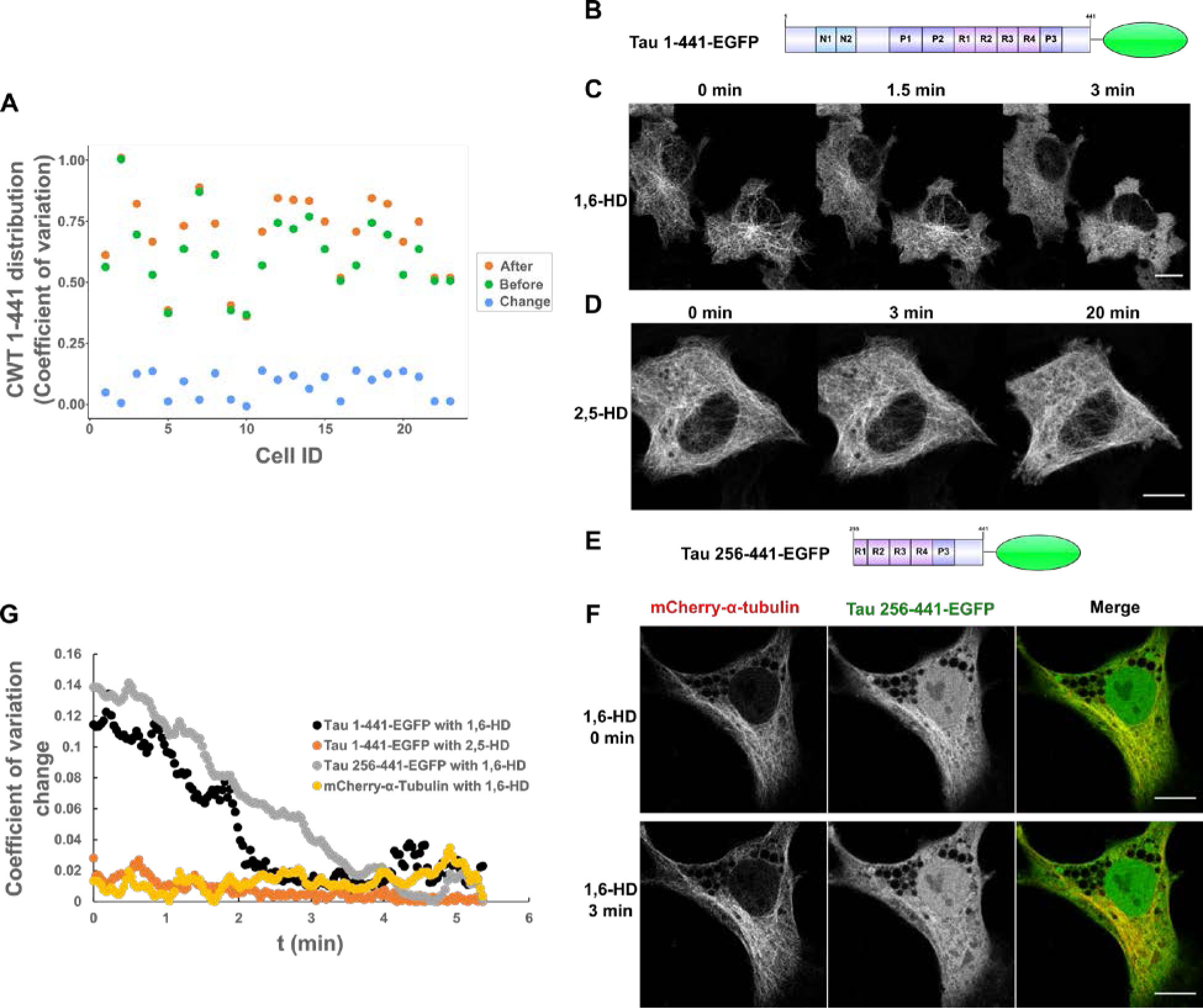
Quantification of Cry2WT-mCherry-tau (CWT 1-441) distribution in SH SY5Y before and after blue light activation and Tau dissociates from MTs after 1,6-hexanediol (1,6-HD) treatment in Hela cells. A) Cry2WT-mCherry-tau (CWT 1-441) distribution in SH SY5Y before activation (green dots), after activation (orange dots) and change by activation (blue dots) were quantified using pixel coefficient of variation (y-axis) for each cell (x-axis). The coefficient of variation of the tau signal after blue light activation is significantly higher than before, n = 23, p = 7.0 × 10^−7^, paired t-test. B-D) Full length tau 1-441-EGFP is sensitive to 1,6-HD but not to 2,5-hexanediol (2,5-HD). B) Protein schematic diagram of tau 1-441-EGFP. C) Tau 1-441-EGFP expressed in Hela cells was treated with 1,6-HD which resulted in the loss of a MT binding pattern to a diffuse pattern. Selected images at 0 min, 1.5 min and 3 min are presented. D) Tau 1-441-EGFP was treated with 2,5-HD in Hela cell, selected images at 0 min, 3 min and extended time to 20 min are presented. Tau binds to MT even after extended treatment of 2,5-HD. E-F) Tau truncations containing MTBD and the extreme C-terminus remain sensitive to 1,6-HD treatment, and MT was not affected. E) Protein schematic diagram of tau 256-441-EGFP from Dr. Gerold Schmitt-Ulms (Gunawardana et al. 2015). F) Tau 256-441-EGFP (green) and mCherry-α-tubulin (red) were overexpressed in Hela cells, treated with 1,6-HD. Montage of selected images at 0 min and 3 min treatment are presented. From left to right, mCherry-α-tubulin, tau 256-441-EGFP and the merge. While truncated tau became diffuse after 3 min, mCherry-α-tubulin still maintained the MT pattern. Note that there is a nuclear population of EGFP signal for tau 256-441-EGFP overexpression, which is commonly seen for EGFP fused with proteins of low molecular weight. G) Quantification of Tau distribution upon 1,6-HD or 2,5-HD treatment based upon coefficient of variation. Distribution of Tau 1-441-EGFP treated with 1,6-HD (black), Tau 1-441-EGFP treated with 2,5-HD (orange), Tau 256-441-EGFP treated with 1,6-HD (grey), and mCherry-a-Tubulin treated with 1,6-HD (yellow) were evaluated and plotted over the time.

**Figure 4-figure supplement 1:**
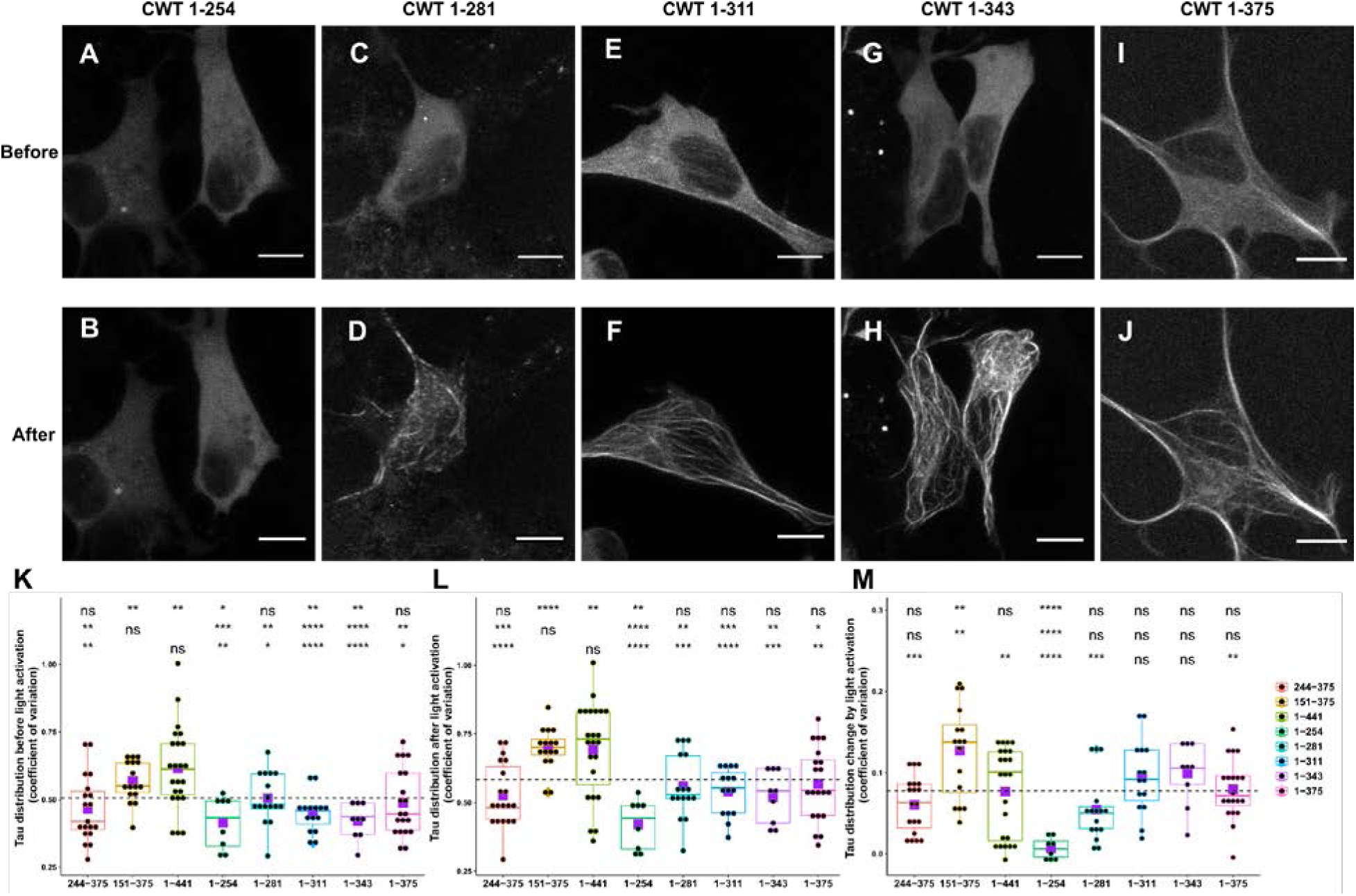
Comparison of Cry2WT-mCherry-tau distribution before and after light activation for various tau constructs. A-J) Cry2WT-mCherry fused various tau constructs were overexpressed in SH SY5Y cells. Signal distribution was studied before and after blue light. Representative maximum intensity Z-projection images before and after blue light of each construct are shown, CWT 1-254 before (A) and after (B), CWT 1-281 before (C) and after (D), CWT 1-311 before (E) and after (F), CWT 1-343 before (G) and after (H), CWT 1-375 before (I) and after (J). Scale Bar, 10 µm. K-M) Quantification of Cry2WT-mCherry-tau distribution before activation (K), after activation (L), and change by activation (M) are shown in boxplot. Constructs from left to right along the x-axis are CWT 244-375, CWT 151-375, CWT 1-441, CWT 1-254, CWT 1-281, CWT 1-311, CWT 1-343, CWT 1-375. Distribution of Cry2WT-mCherry-tau is evaluated by pixel coefficient of variation over the cell shown along the y-axis. n = 19, 16, 23, 8, 16, 15, 9, 20, for constructs from left to right along the x-axis. Mean for each group is indicated by a purple square, also listed in Figure 4-table supplement. Base-mean for all the constructs as a whole is indicated as a horizontal dot line. Statistical significance by pairwise t-test of each independent group is indicated at the top of each plot, from top to bottom, against all the constructs as a whole, against CWT 1-441 and against CWT 151-375, correspondingly. ns, not significant; *, p < 0.05; ** p < 0.01; ***, P < 0.001; ****, P < 0.0001.

**Figure 4-table supplement:** Summary of the mean and standard error (S.E.) for each Cry2WT-mCherry-Tau construct, in cases of before activation, after activation, and change-by-activation.

| | Mean ± S.E. | | |
| --- | --- | --- | --- |
| **CWT construct** | **Before activation** | **After activation** | **Change by activation** |
| **244-375** | 0.465 ± 0.028 | 0.526 ± 0.027 | 0.0606 ± 0.008 |
| **151-375** | 0.568 ± 0.019 | 0.695 ± 0.020 | 0.127 ± 0.014 |
| **1-441** | 0.616 ± 0.032 | 0.693 ± 0.036 | 0.077 ± 0.011 |
| **1-254** | 0.414 ± 0.035 | 0.42 ± 0.032 | 0.00664 ± 0.005 |
| **1-281** | 0.505 ± 0.024 | 0.56 ± 0.029 | 0.0548 ± 0.010 |
| **1-311** | 0.446 ± 0.018 | 0.54 ± 0.022 | 0.0944 ± 0.012 |
| **1-343** | 0.421 ± 0.022 | 0.52 ± 0.032 | 0.0992 ± 0.013 |
| **1-375** | 0.489 ± 0.028 | 0.568 ± 0.030 | 0.0798 ± 0.008 |
| **Base-Mean** | 0.505 ± 0.011 | 0.583 ± 0.012 | 0.078 ± 0.077 |

**Figure 5-figure supplement 1:**
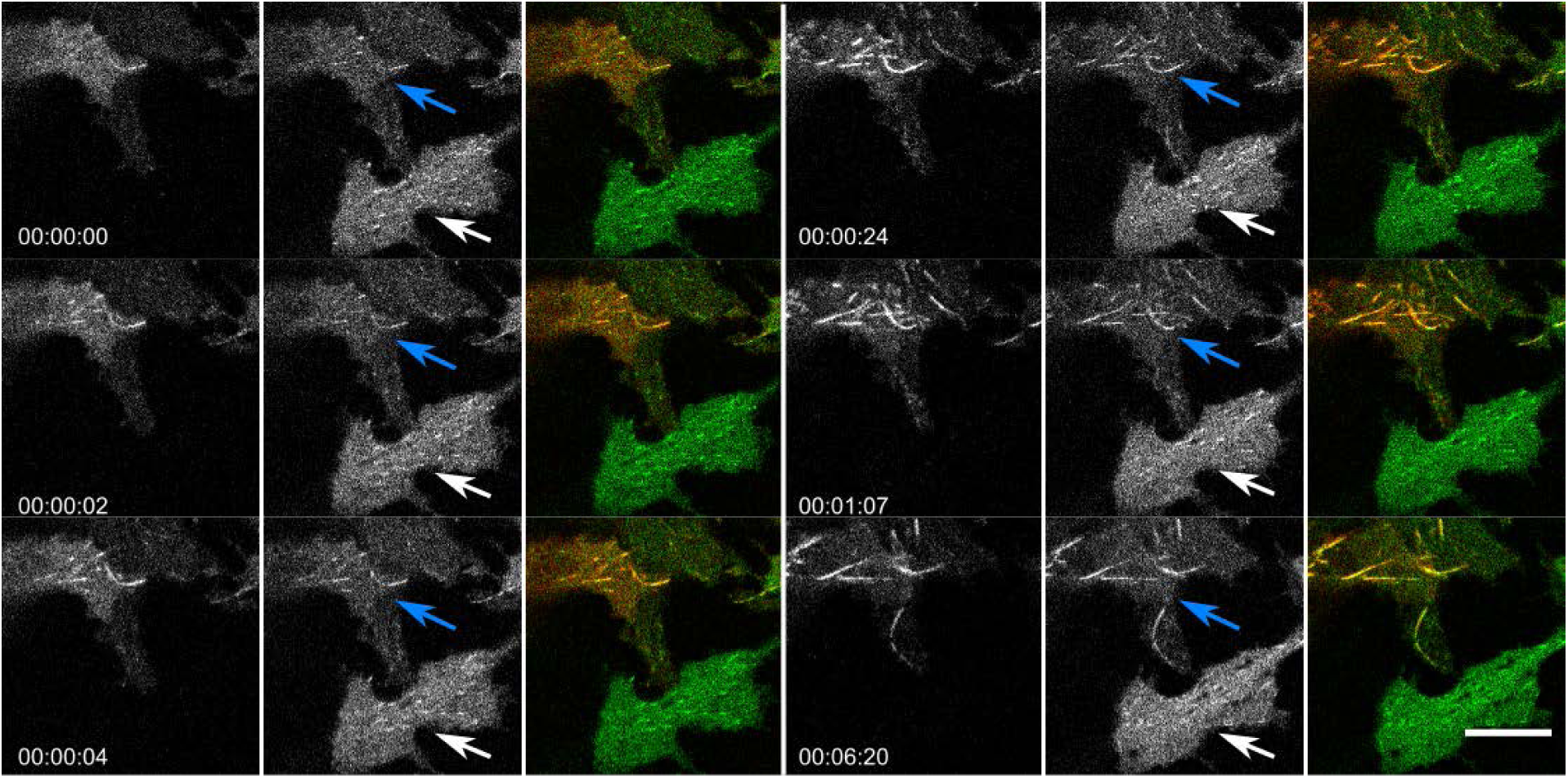
Cry2WT-mCherry-tau 151-254 (CWT 151-254) /EB1-EGFP bound along the MT lattice of MT bundles upon light activation. Time lapse of EB1-EGFP (green) when overexpressed with Cry2WT-mCherry-tau 151-254 (CWT 151-254, red) in SH SY5Y cells activated by shallow blue light (488 nm, 5% power). Note that besides binding MT +TIPs and moving as comets, upon light activation, some CWT 151-254/EB1-EGFP extended the binding along the MT lattice and MT bundles (following the blue arrows in the time lapse) while the cell at the bottom which also has expression of EB1-EGFP but no CWT 151-254 does not show shift of EB1 localization from MT +TIPS, remaining as comet (white arrows). Montage of two fluorescent channels is presented in the order from left, the mCherry channel, the GFP channel separate and the merged. Scale Bar, 10 µm.

## Supplemental Information

### Supplemental Method of FTS-Figure 6C

Recently we developed a coarse-grained polyampholyte model for studying charge-sequence dependent phase behavior of tau in the presence of RNA (Lin et al. 2019). A numerical method from the polymer physics community known as field theoretic simulation (FTS) was introduced to overcome the limitations of traditional molecular dynaimcs simulation. The method begins by transforming the canonical partition function of the coarse-grained model into a complex-valued statistical field theory by an exact Hubbard-Stratonovich transformation.

The fluctuating chemical and electrostatic fields are sampled using complex Langevin (CL) simulation. The fields were sampled on a spatial grid of 34^3^ sites. An exponential time difference (ETD) algorithm was used to numerically propagate the CL equations of motion with a timestep of 0.02. All simulations were performed in a cubic box of length 5.55 nm using periodic boundary conditions.

In this method, each amino acid is represented by a single interaction site of size b. All microscopic site densities are smeared over a finite volume by convolution with a Gaussian profile. The segment size b determines the length scale of the system. To parameterize our model we set b = 0.4 nm which approximates the distance between *C_α_* carbons in the protein backbone. Charged residues i and j interact via the electrostatic potential 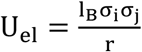, where σ_i_ is the charge on residue i at pH 7.4. The parameter l_B_ is the Bjerrum length defined as 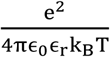 with e the unit of electronic charge, ϵ_0_ the vacuum permittivity, and ϵ_r_ the relative dielectric strength of the solvent. To parameterize the model, we set *ϵ_r_* = 80 for water. In our model, l_B_ is an input parameter, which characterizes the strength of the electrostatic interaction energy. For reference, at 300 K, for water, l_B_ = 0.71 nm. By running multiple simulations each at different values of l_B_, we can determine the phase coexistence points as a function of l_B_ and compare how changes in the primary amino acid sequence widens or narrows the window for coacervate through charge-sequence effects. Besides Coulomb interactions, the polymer segments in implicit water interact via a short-ranged excluded volume repulsion potential of strength υ. To compare among tau constructs, we include a negatively charged polyelectrolyte species with the same degree of polymerization in order to model co-coacervating species such as single-stranded RNA. For tau 255-441 and tau 151-254 regions, the volume fraction was chosen to maintain a 1:1 charge ratio between polyampholyte and polyanion species. For the N-terminus tau 1-150 and C-terminus tau 376-441 sequences, which are net negatively charged, total charge neutrality was maintained by adding positively charged sodium ions to balance the net negative charge from the polymer species.

To compare the droplet stability of the PRD (151-254) with the N-terminus (1-150) region, we perform FTS simulations in the Gibbs ensemble (Riggleman and Fredrickson 2010). In these simulations a dense (coacervate) phase and a dilute (solution) phase are simultaneously simulated in separate cells. Together these simulation cells form a canonical ensemble for which both the total mass and total volume is conserved. Chemical and mechanical equilibrium is achieved by performing mass and volume swaps between the two simulations cells while imposing charge neutrality constraints. The concentration and relative box volumes are updated periodically until convergence is achieved. The results of a Gibbs ensemble simulation performed with l_B_ = 0.71 nm and 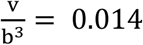, which corresponds roughly to T = 293 K in our model, shows that the PRD has a dilute phase and concentrated (droplet) phase that reach a stable coexistence, while the simulation cells for the N-terminus region converges to a single phase of homogenous bulk density, corresponding to the case in which the droplet vanishes.

